# Structure of nucleotide-bound Tel1^ATM^ reveals the molecular basis of inhibition and structural rationale for disease mutations

**DOI:** 10.1101/696203

**Authors:** Luke A. Yates, Rhys M. Williams, Sarem Hailemariam, Rafael Ayala, Peter Burgers, Xiaodong Zhang

## Abstract

**SUMMARY:** Yeast Tel1 and its highly conserved human orthologue ATM are large protein kinases central to the maintenance of genome integrity. Mutations in ATM are found in ataxia-telangiectasia (A-T) patients and ATM is one of the most frequently mutated genes in many cancers. Using cryo electron microscopy, we present the structure of Tel1 in a nucleotide-bound state. Our structure reveals molecular details of key residues surrounding the nucleotide binding site and provides a structural and molecular basis for its intrinsically low basal activity. We show that the catalytic residues are in a productive conformation for catalysis, but the PIKK-regulatory domain-Insert (PRD-I) restricts peptide-substrate access and the N-lobe is in an open conformation, thus explaining the requirement for Tel1 activation. Structural comparisons with other PIKKs suggest a conserved and common allosteric activation mechanism. Our work also provides a structural rationale for many mutations found in A-T and cancer.

## INTRODUCTION

*Saccharomyces cerevisiae* Tel1 and its highly conserved human orthologue Ataxia-Telangiectasia-Mutated (ATM), are major kinases responsible for maintaining genome integrity and is highly conserved in Eukaryotes. Tel1^ATM^ is recruited to sites of DNA damage by the MRX/N complex, a key element in double-strand (ds)DNA break repair comprising of <u>M</u>re11, <u>R</u>ad50, and <u>X</u>rs2 (<u>N</u>bs1 in human) (Falck et al., 2005). MRX/N is also the major activator of Tel1^ATM^, which subsequently phosphorylates hundreds of targets that contain a S/T-Q motif (Lee and Paull, 2004; Matsuoka et al., 2007). Many of these targets, including CHK1/2, ATM, BRCA1, PALB2, p53 and H2AX, are tumour suppressors involved in cell cycle control and dsDNA break repair through homologous recombination (Lavin and Kozlov, 2007; Matsuoka et al., 2007). The precise mechanism of Tel1^ATM^ activation is not fully understood, but is suggested to involve autophosphorylation (Kozlov et al., 2006), lysine acetylation (Sun et al., 2005) and dissociation of homodimers into monomers (Bakkenist and Kastan, 2003). Critically, mutations in ATM are found in Ataxia-Telangiectasia (A-T), a rare disease primarily associated with immunodeficiency and progressive neurological decline. A-T patients also have an increased susceptibility to malignancy due to genomic instability, and ATM is one of the most frequently mutated genes in many cancers (Choi et al., 2016).

Tel1^ATM^ belongs to a family of PI-3K (phosphatidylinositol-3-kinase)-like kinases that also includes ATR (ATM-Rad3-related, and its yeast orthologue Mec1), DNA-PKc, mTOR (mammalian target of Rapamycin), TRRAP/Tra1 and SMG1 (Baretić and Williams, 2014). All PI-3K kinases contain a canonical two-lobed kinase domain, with the smaller N-lobe containing the highly conserved Glycine-rich loop (Gly-rich loop), whilst the larger C-lobe possesses the catalytic and activation loops (Walker et al., 1999). A number of additional conserved and functional elements within the kinase domain have been identified in PIKKs including the LBE (LST8-binding element) (Yang et al., 2013) and the PRD (PIKK regulatory domain) (Mordes et al., 2008). Flanking the kinase domain are the N-terminal HEAT repeats, followed by a FAT (<u>F</u>RAP [FKBP12-rapamycin associated protein], <u>A</u>TM, <u>T</u>RRAP [transformation/transcription domain associated protein]) domain and a ∼35 residue FATC domain at the C-terminus (Imseng et al., 2018). Due to their large sizes, structural and mechanistic studies on these kinases have been challenging. A number of high-resolution structures (< 4.0 Å, where many side chains can be resolved) are available, mostly of those involving mTOR, including its complexes with a number of activator proteins (Yang et al., 2017). The 4.3 Å crystal structure of DNA-PKcs revealed the architecture of this large PIKK (Sibanda et al., 2017) whereas the 3.9 Å cryoEM structure of Mec1-Ddc2 provided a structural basis for how ATR^Mec1^ might be kept in an inhibited state (Wang et al., 2017). The cryoEM structures of Tel1 (Xin et al., 2019) and human ATM (Baretić et al., 2017) in the absence of nucleotides have shed some light into kinase function but do not provide a complete understanding of the role of the regulatory elements in maintaining an auto-inhibited state.

Here we present the cryoEM structure of *Saccharomyces cerevisiae* Tel1 in complex with an ATP analogue, AMP-PNP, with sufficient resolution (3.7 – 4.0 Å) to resolve the bound nucleotide and allow most of the side chains in the conserved FAT-Kinase-FATC (FAT-KIN) to be resolved (**Figure 1, Figure S1 and S2**). The structure reveals that the active site residues of Tel1 are in a productive conformation with the exception that the N-lobe, which also contributes to catalytic activities, is likely to undergo further closure upon activation. Significantly, a conserved insertion in the PRD (PRD-I) obscures peptide substrate access to the active site. This explains the low intrinsic activity of Tel1 and its requirement for activation by the binding of MRX and DNA. Comparisons with mTORC1 and DNA-PK structures suggest a common allosteric activation mechanism that could lead to the closure of N-lobe as well as the relocation of the PRD-I domain, leading to an active site that is fully accessible for peptide substrate binding. Due to the high degree of structural conservation between ATM and the Tel1 FAT-KIN regions, where a large number of pathogenic mutations are located, our structure also provides a structural basis for many disease mutations found in ATM.

**Figure 1.**
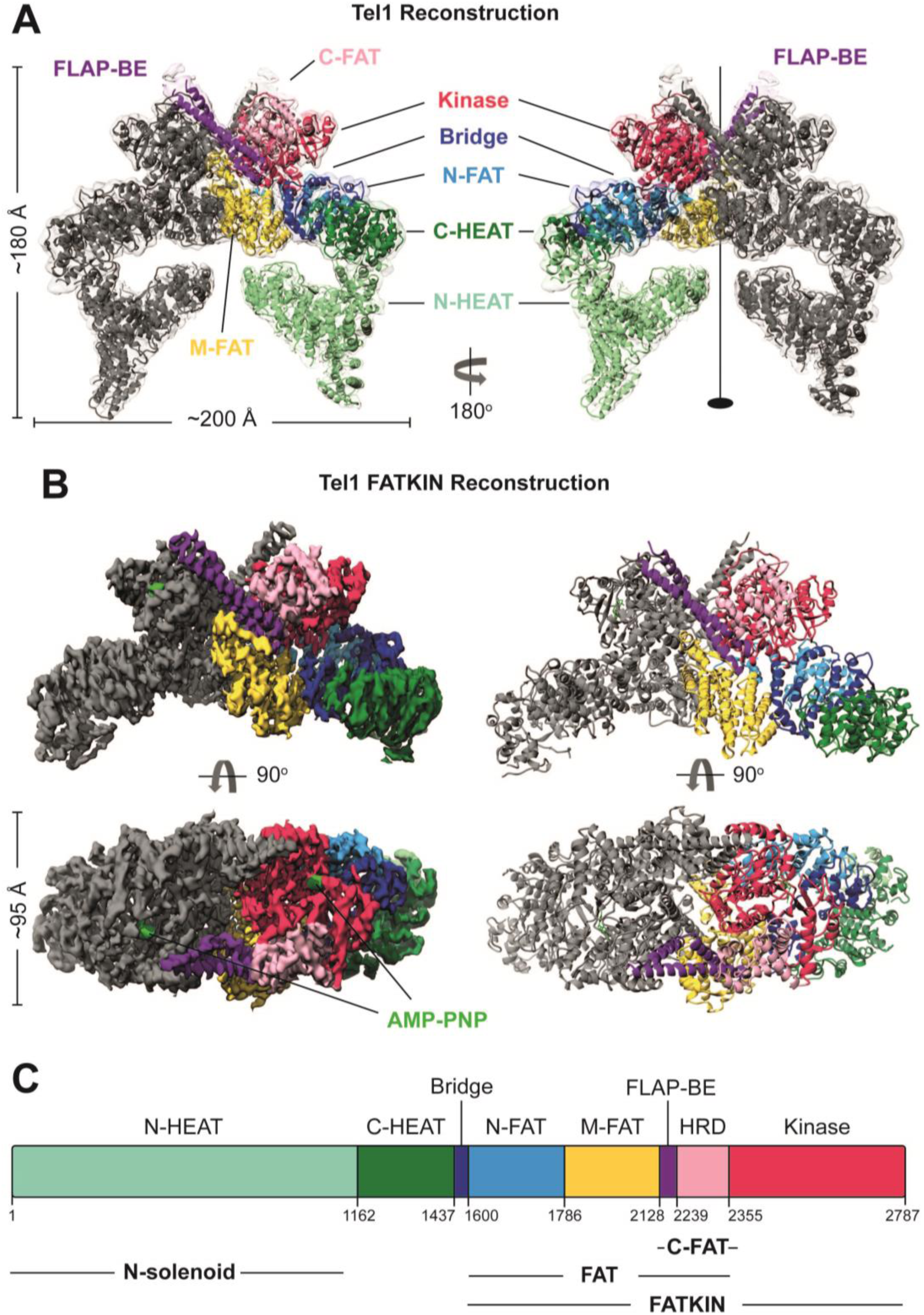
Structure of Tel1 Dimer. (A) Overall structure of the nucleotide bound (AMP-PNP) Tel1 dimer structure determined by cryoEM to 4.0 Å resolution by Gold Standard Fourier shell correlation (GS-FSC). Regions of the structure are coloured according to domains; Kinase, red; C-FAT, pink; FLAP-BE, purple; N-FAT, blue; Bridge, dark blue; C-HEAT, green; N-HEAT, light green. Approximate dimensions of the protein are also given. (B) Reconstruction of the FATKIN region determined to 3.7 Å resolution by GS-FSC, coloured as in (A). (C) Domain arrangement of Tel1.

## RESULTS AND DISCUSSION

### Structure of the Tel1 dimer

Tel1 was expressed and purified as reported previously (Sawicka et al., 2016) and its kinase activity shown to be stimulated synergistically by both double-stranded (ds) DNA and MRX (Hailemariam et al., 2019). The overall Tel1 dimer structure was refined to a global resolution of 4.0 Å (**Figure S2**). Using localised masking around density regions corresponding to the FAT-KIN domains as well as regions of HEAT repeats, an improved resolution of 3.7 Å was obtained for these regions, which correspond to two thirds of the whole protein. In key functional regions around the kinase domain and the dimer interface, the local resolution estimates are better than 3.5 Å; and densities for bulky side-chains are clearly visible (**Figure S2**). Consequently, we could build side chains into the majority of the FAT, kinase and FATC domains (residues 1527-2787) and place confident sequence assignment between residues 968 and 1526. The N-terminal HEAT repeats are flexible, as was also shown in ATM structures, and left as Cα trace only due to the lack of clear side chain density. Our structure is in excellent agreement with that of an ATM closed dimer, which was limited to Cα-Cβ atoms (Baretić et al., 2017). Interestingly, we did not observe an open dimer conformation or monomers in our analysis.

Different subdomain terminologies have been used to describe PIKKs, including ATM. For simplicity, we use rigid bodies to define subdomains within the HEAT repeats and the FAT domain, which was originally assigned by multiple sequence alignments (Bosotti et al., 2000). Specifically, by comparing the structure of Tel1 and those of ATM and mTOR, the N-terminal HEAT repeats can be separated into two rigid bodies, which we term N-HEAT (residues 1-1162) and C-HEAT (residues 1163-1437) (**Figure 1**). There is a connecting region, denoted a bridge domain (residues 1438-1600) before the FAT domain, which we have extended and also divided into several rigid bodies: N-FAT (N-terminal FAT, 1600-1786), M-FAT (Middle-FAT, 1787-2128) and C-FAT (C-terminal FAT, 2129 -2355) (**Figure 1 and Figure S3**). The C-FAT consists of the FLAP-BE (as defined in (Baretić et al., 2017)) and HRD domain (**Figure 1**).

The Tel1 dimer interface is extensive and can be divided into three layers (**Figure 2A**); the top layer consists of PRD-I and LBE of one protomer interacting with a long helical antenna specific to Tel1^ATM^ called FLAP-BE, of the adjacent protomer (FLAP-BE’, where ‘denotes elements from the adjacent protomer) (**Figure 2**); the middle layer mainly consists of FATC of one protomer and the adjacent M-FAT’ (immediately preceding FLAP-BE) of the other protomer, while the bottom layer consists of M-FAT-M-FAT’ interactions **(Figure 2A**). The buried surface area at the dimer interface is extensive with over 3000 Å^2^ per protomer. The bottom layer of the dimer interface is exclusively hydrophobic in nature, with a vast array of large bulky hydrophobic residues (Phe and Leu) forming a well-buried core around the 2-fold symmetry axis (**Figure 2B**). The middle layer employs a mixture of polar and hydrophobic residues (**Figure 2C**), whereas the upper layer, which has the smallest interface surface area and is the least well resolved, interacts via charge complementarity (**Figure 2D**). Given the characteristics of the dimer interface, it seems that a dimer-to-monomer transition would be energetically unfavourable. Furthermore, the hydrophobic nature of the bottom layer enables some conformational flexibility at the bottom layer of the dimer interface, whereas the top layer, which is polar in nature, can be broken or rearranged. Indeed, a relative rotation of the FAT-KIN regions in the ATM dimer has been observed in the open dimer conformation (Baretić et al., 2017).

**Figure 2.**
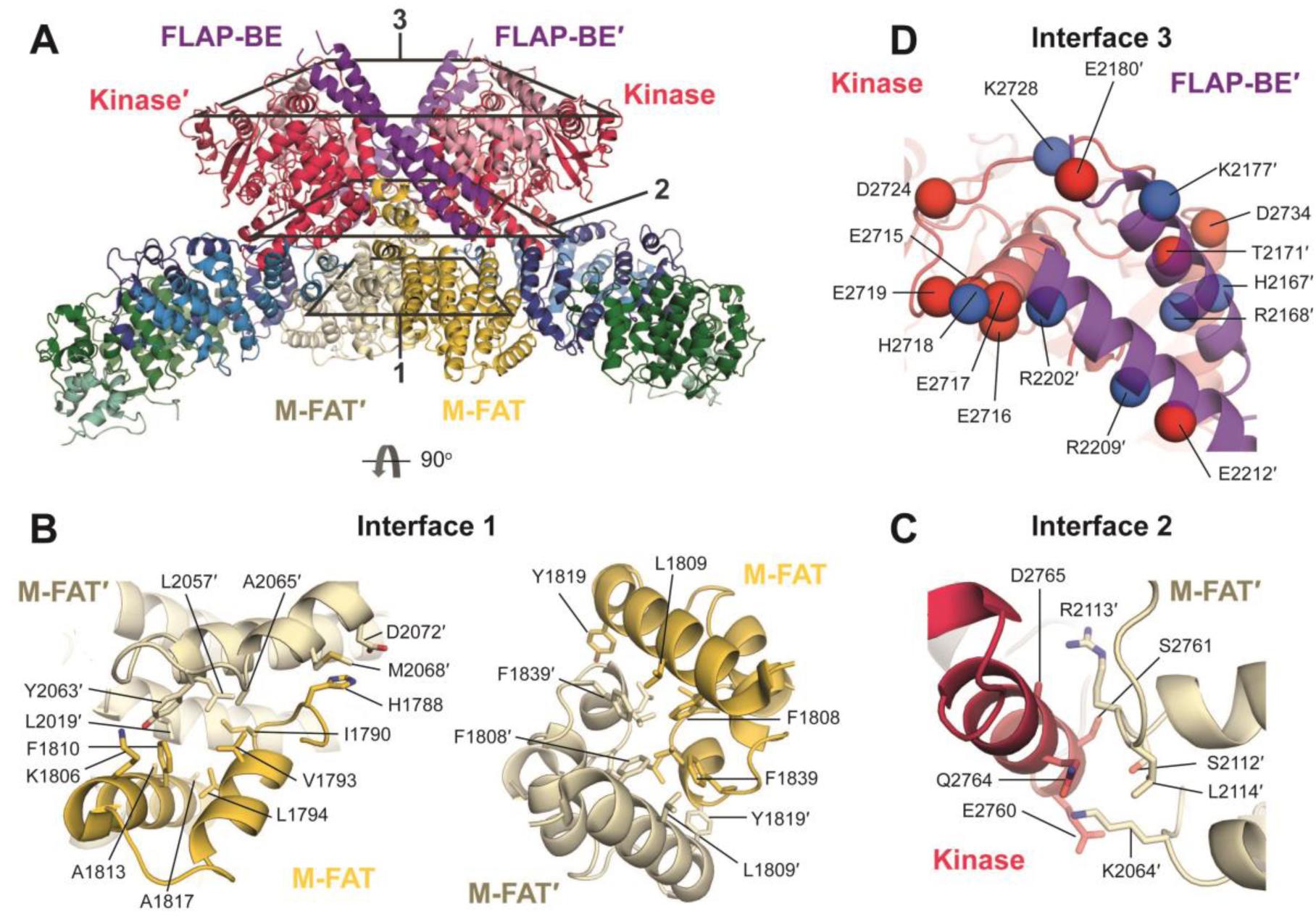
Tel1 Dimer Interface. (A) FATKIN dimer structure coloured by domain as in Fig1, and for clarity M-FAT’ is coloured in pale yellow to distinguish it clearly. Different layers of the dimer interface are boxed and detailed views of the nature of the interface are shown in (B-D).

### Tel1 nucleotide-bound kinase domain is in a catalytically productive conformation

Similar to the other PIKKs such as mTOR, ATM, DNA-PKc and Mec1, the kinase domain is surrounded by the FAT domains that form a C-shaped cradle surrounding the kinase domain (**Figure 3A**). It is worth noting that N-FAT contacts the kinase C-lobe whereas C-FAT contacts the N-lobe (**Figure 3A, Figure 1A**). M-FAT sits away from the kinase domain and instead contacts C-HEAT and is in close proximity with N-HEAT and could therefore transmit conformational changes from the HEAT region to the kinase domain (**Figure 3A, Figure 1A**).

**Figure 3.**
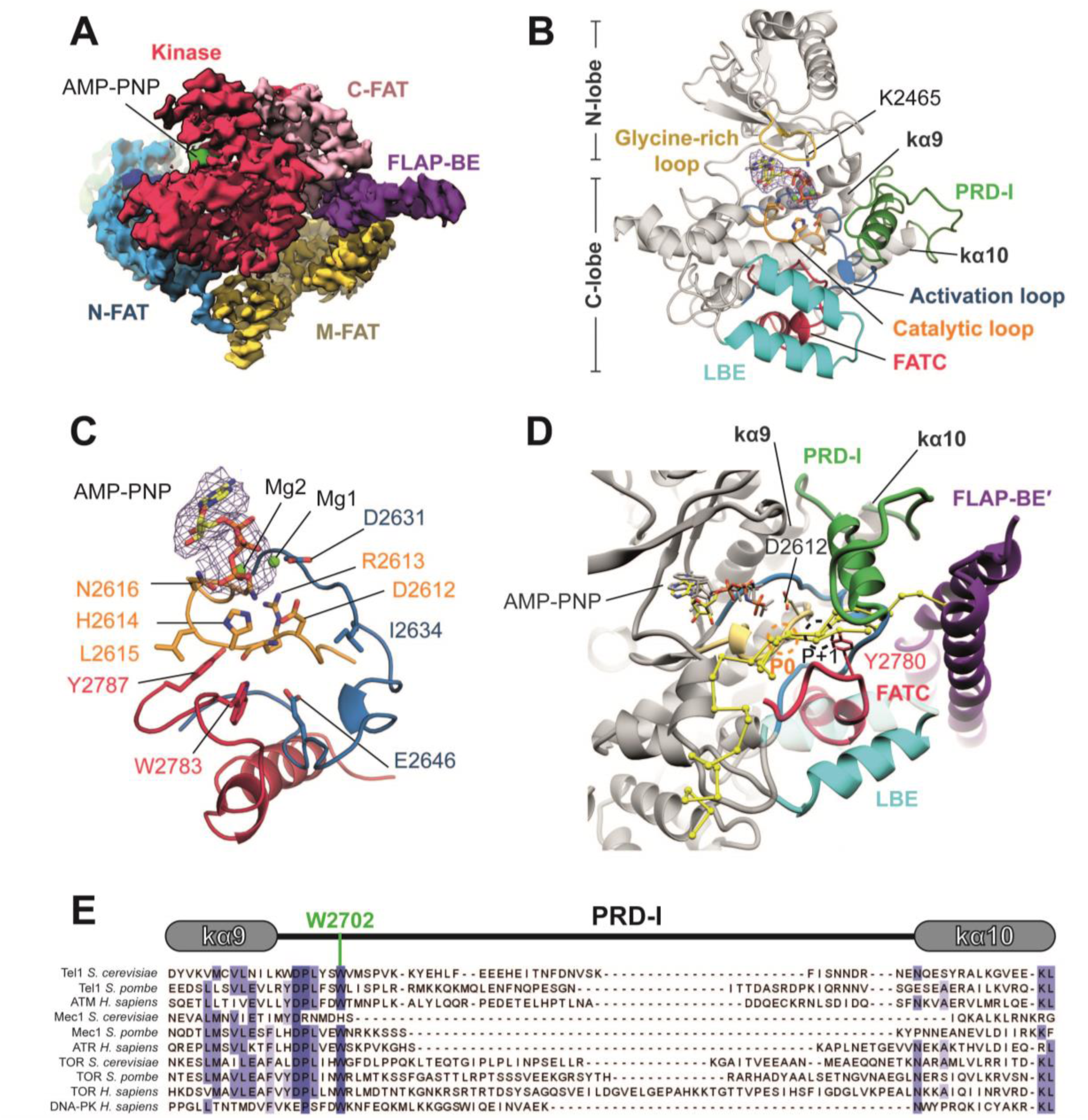
Details of the Tel1 Kinase Domain and the PRD-I. (A) CryoEM density of Tel1 FAT-KIN shows the FAT domains cradle the kinase domain. Domains are coloured as in Fig1. (B) Structural details of the AMP-PNP bound (with cryo EM density shown as mesh) kinase domain (grey) with key structural features highlighted and labelled. (C) Molecular details of interacting residues with AMP-PNP, the activation loop (blue) and FATC (red) which support the catalytic loop (yellow). (D) Predicted peptide binding using superimposition of kinase-substrate peptide complex crystal structures of PKA, Pak-4 and Cdk2. The target serine/threonine is shown and labelled (P0) and the predicted location of the P+1 residue is highlighted and is close to Y2780 of the FATC. (E) Sequence alignment of the structurally conserved kα9 and kα10 and the intervening PRD of several PIKKs. An invariable tryptophan (W2702) that anchors the PRD is highlighted.

The structure of nucleotide-bound Tel1 shows the catalytic residues of the C-lobe in a catalytically productive conformation, with clear density for AMP-PNP **(Figure 3B-C)**. The catalytic loop, containing the highly conserved ^2612^DRH^2614^ motif, adopts a similar conformation to that observed in mTOR structures **(Figure S4)** (Yang et al., 2013, 2017), where D2612 is well-positioned to act as the catalytic base for deprotonation of the hydroxyl group from the peptide substrate, and the side chains of H2614 and N2616 co-ordinate the γ-phosphate of AMP-PNP (**Figure 3B-C**) (Kenyon et al., 2012). The Mg^2+^-binding ^2631^DLG^2633^ motif and the adjacent activation loop are well-ordered, allowing us to confidently model in two Mg^2+^ ions co-ordinating the β-/γ- (Mg1) and α-/γ-phosphates (Mg2) of AMP-PNP into the density, using nucleotide-bound structures of PKA and mTORC1 as a guide (**Figure 3B**) (Das et al., 2015; Yang et al., 2013). The activation loop, which is ordered in this kinase, wraps around the catalytic loop and supports kinase activity via several interactions. Most notably, residues E2646 and I2634 of the activation loop sandwich the catalytic loop, and there is an apparent interaction between R2613 of the catalytic loop (^2612^DRH^2614^ motif) and the main chain of the activation loop, close to the ^2631^DLG^2633^ motif. The catalytic loop is further constrained by two aromatic residues (W2783 and Y2787) from the C-terminus of FATC (**Figure 3C**). These structural constraints explain the highly conserved nature of FATC and why mutations in this region abolish Tel1^ATM^ activity (Jiang et al., 2006).

The binding pocket for the adenosine ring of AMP-PNP is formed by a number of hydrophobic residues from both N-and C-lobes of the Tel1 kinase domain, with the glycine-rich loop from the N-lobe forming a partial lid over the bound nucleotide (**Figure 3B**). The highly conserved residue K2465 from the N-lobe contributes to α-/β-phosphate coordination of the bound AMP-PNP and is equivalent to K72 of PKA, which is essential for catalysis (Iyer et al., 2005) (**Figure 3B**). Interestingly, comparison of the Tel1 active site with nucleotide-bound mTOR in non-activated and RHEB-activated states (Yang et al., 2017) shows that the N-lobe and the glycine-rich loop of Tel1 most closely resemble that of non-activated mTOR (**Figure S4**). Further closure of the mTOR N-lobe and glycine-rich loop occur upon activation. It is therefore likely that the N-lobe and glycine-rich loop of Tel1 will also undergo further closure as part of the catalytic cycle. In addition to the glycine-rich loop, the active site is further surrounded by LBE, PRD and FLAP-BE’ (**Figure 3**), highlighting the importance of these elements in regulating active site access.

### Tel1^ATM^ PRD-I and FATC regulate substrate binding

In addition to the open N-lobe, our structure shows that the active site cleft of the Tel1 kinase domain is restricted by the PIKK-regulatory domain (PRD), which we have denoted as the PRD-Insert (PRD) (see below, Figure 3B). The PRD is a PIKK-specific feature that was originally identified in ATR as an element between the kinase domain and the FATC that is essential for the binding of the activator TopBP1 (Mordes et al., 2008). Subsequent PIKK structures have revealed that the regions proposed to be involved in TopBP1 binding are in the highly conserved Kα10 helix, whereas the regions between Kα9 and Kα10 are variable in length and composition between different PIKKs **(Figure S5)**. Therefore we denote this variable region as the PRD-Insert (PRD-I), which is equivalent to Kα9b, Kα9c and Kα9d identified in mTOR and ATM structures (Baretić et al., 2017; Yang et al., 2013, 2017).

In the Tel1 and ATM closed dimer structures, the PRD-I, although only partly resolved, is predicted to occupy the position of a peptide substrate (Baretić et al., 2017; Xin et al., 2019). In the nucleotide-bound Tel1 structure, the complete PRD-I could be resolved. It structurally connects Kα9-Kα10 at the back of the kinase domain to the LBE and FATC at the front of the kinase domain (**Figure 3B**). Starting from Kα9, the PRD-I displays an extended loop followed by a single α-helix, situated directly above the activation loop, reaching towards LBE and FLAP-BE’ before returning to Kα10 via a loop (**Figure 3B**). Importantly, PRD-I interacts with many of the key functional and structural elements, including the activation loop, the FATC, the LBE and the FLAP-BE’ (**Figure 3B and 3D**). At its N-terminus, an invariant W2702 of PRD-I inserts into the hydrophobic pocket between Kα9 and Kα10, (**Figure 3B**). Towards the C-terminal end, acidic residues in PRD-I (^2715^EEEHE^2719^) are in close proximity to a positively charged patch in FLAP-BE’ (^2197^KRHYHR^2202^) (**Figure 2D**). Together these elements occlude the access to the active site of Tel1.

To estimate the path of a bound Tel1 peptide substrate, we aligned the Tel1 kinase domain with the catalytic and activation loops of substrate-bound structures of PKA (PDB: 3×2u) (Das et al., 2015), Cdk2 (PDB: 3qhw) (Bao et al., 2011) and Pak-4 (PDB: 4jdh) (Chen et al., 2014) **(Figure 3D)**. It is clear that PRD-I in Tel1 overlaps with the peptide substrate-binding site, whereas the γ-phosphate of AMP-PNP and the catalytic aspartate (D2612) are suitably close to the predicted location of the target Ser/Thr (P0) for phosphotransfer to occur (Das et al., 2015) (**Figure 3D**). The PRD-I, together with the LBE and FLAP-BE’, thus occludes the active site and occupies substrate binding site. It is therefore the major determinant for inhibition and the requirement of Tel1 activation by MRX-dsDNA. Furthermore, Y2780 within the FATC sits close to the predicted P+1 peptide substrate position, suggesting a potential role in determining substrate specificity. This is consistent with a Y2780A mutant Tel1 demonstrating significantly defective levels of Rad53 phosphorylation, supporting the idea that this part of FATC may be involved in target recognition (Ogi et al., 2015). We thus suggest a dual role for FATC, being important for constraining the catalytic loop conformation as well as substrate peptide recognition.

### Proposed activation mechanism of Tel1^ATM^

Among the PIKK enzymes, only mTOR has a set of high-resolution structures that enable side chain assignments for both non-active and activated states. Comparisons of mTORC1 and the activator bound mTORC1-RHEB complex reveal that RHEB binding induces allosteric conformational changes to C-FAT that, via a clockwise rotation around the kinase C-lobe, results in the closure of the N-lobe and glycine-rich loop, thus realigning the active site (Yang et al., 2017). RHEB binds at a significant distance away from the kinase domain, close to N-FAT (Yang et al., 2017). Upon binding of RHEB there is little conformational change in N-FAT. Instead, through HEAT domain relocation, significant conformational changes occur in M-FAT and are subsequently transmitted to C-FAT (Yang et al., 2017). DNA-PKc also binds to its activator, Ku70-Ku80-DNA, through its HEAT repeats and structural comparisons between mTOR and DNA-PKc alone and in complex with their activators reveal a common mode of allosteric activation involving domain movement of M-FAT and C-FAT (Sharif et al., 2017; Sibanda et al., 2017; Yin et al., 2017). Despite the structural and sequence divergence of the N-HEAT region, the FAT-KIN domains of PIKKs are highly conserved in their structures (**Figure S6**). Furthermore, the conserved conformational changes in mTOR and DNA-PKc, irrespective of their distinct activator binding sites, suggest similar conformational changes might occur in Tel1 as a means for activation.

Intact MRX/MRN complexes along with dsDNA are required to stimulate Tel1^ATM^ activity (Hailemariam et al., 2019; Lee and Paull, 2004, 2005; Lee et al., 2003; Uziel et al., 2003). Tel1^ATM^ interacts with MRX/N via direct associations with the complex (Lee and Paull, 2004), and recent studies indicate that each of the three MRX subunits binds Tel1, suggesting an extensive MRX-Tel1 interaction interface (Hailemariam et al., 2019). The interaction between Tel1 and the C-terminus of Xrs2/Nbs1, has been mapped to regions in the N-HEAT and C-HEAT domains in Tel1^ATM^ (You et al., 2005) in a similar location to DNA-PK and mTORC1 activator binding sites (**Figure S6**). When we align the Tel1 and the inactive mTORC1 structures on their kinase C-lobes, N-FAT and C-FAT align reasonably well, with the largest differences in M-FAT (**Figure S6**). It is therefore plausible that binding of MRN/X would result in a concerted motion of HEAT and M-FAT which is propagated to C-FAT, resulting in the N-lobe and glycine-rich loop closure analogous to RHEB-activated mTORC1 (Yang et al., 2017). Distinct from mTOR, these movements would involve FLAP-BE within the C-FAT, which subsequently could affect PRD-I conformation and the dimer interface, presumably releasing inhibition for substrate binding and active site access. Recent cryoEM structures of apo-Tel1 suggested domain motions which are reminiscent of those observed during mTOR activation, with these intrinsic motions also resulting in increased PRD-I disorder (Xin et al., 2019). Interestingly, we did not observe similar domain movement or PRD-I disorder when Tel1 is nucleotide-bound. This lack of domain movement and a consistently ordered PRD-I maintaining autoinhibition are consistent with observed low-level kinase activity of Tel1 in the absence of activators (∼0.065 phosphates transferred per minute per protomer) (Hailemariam et al., 2019).

### Structural basis for hyperactive mutations of Tel1 and disease associated mutants in ATM

The role of PRD-I in inhibiting the activity of Tel1 and a proposed allosteric activation mechanism are supported by Tel1 hyper-active mutations identified in a genetic screen to rescue Mec1 deficient cells (Baldo et al., 2008). Many mutations are in similar positions to cancer-associated mTOR hyper-activating mutations known to increase sensitivity to allosteric activation (Yang et al., 2017). Several mutations map onto Kα9 and Kα10 and surrounding regions (**Figure 4A**). Among the mutant strains, three possess only single amino acid substitutions (N2692D, F2576V and Q2764H), that hyper-activate Tel1 activity. In one such mutant, N2692 of Kα9 is next to G2252 of C-FAT, which is also mutated in one of the strains, and therefore N2692D or G2252C could affect Kα9 conformation and consequently affect the PRD-I position (**Figure 4B**). Q2764 is within the FATC and also located at the dimer interface with M-FAT’, specifically in a region just before the FLAP-BE’. A Q2672H substitution could affect FLAP-BE’ to aid the relocation of PRD-I. There are also a number of mutations at the interface between N-FAT and C-lobe, close to the LBE and activation loop. These mutations could also affect substrate access (**Figure 4A**). Several mutations, e.g. I2336T and A2287V, are found within the C-FAT, close to the N-lobe and are similar to known mTOR hyper-activating mutations (Yang et al., 2017). Presumably these mutations could allow the N-lobe to move closer to the active site in the absence of activator proteins.

**Figure 4.**
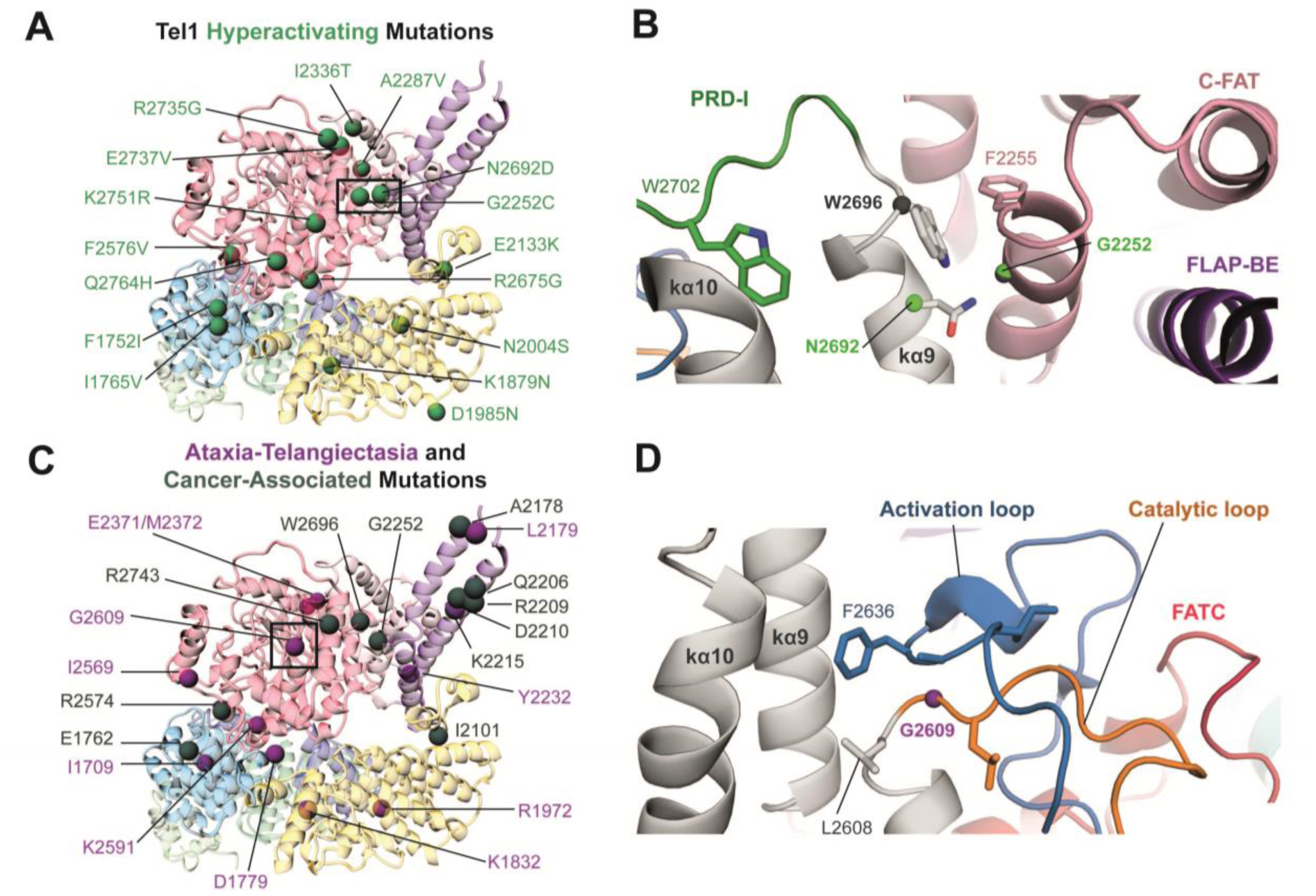
Mapping of Tel1 hyper-activating and ATM disease associated mutations. (A) Tel1 FAT-KIN protomer is shown coloured as in Fig1 with Hyper-activating mutations found in (Baldo et al., 2008) mapped and shown as green spheres. (B) Molecular details of a cluster of mutations in kα9 found in cancer and described in hyper-activating Tel1 phenotypes. (C) Cancer-associated (dark grey spheres) and Ataxia Telangiectasia-associated (A-T, purple spheres) mutations at structurally equivalent locations (see Table S1 for details). (D) Molecular details of G2609 (G2867 in ATM), which is mutated to Arginine in A-T.

The high structural homology between Tel1 and ATM FAT-KIN regions allows us to map pathogenic A-T missense and cancer-associated mutations onto the structure (**Figure 4C**). Ataxia telangiectasia is caused by bi-allelic mutation of the *ATM* gene and a significant proportion of A-T mutations result in abolished ATM expression, via truncation, incorrect splicing or incorrect frame-shifts (McKusick, 2007; Sandoval et al., 1999; Stankovic et al., 1998), suggesting that the molecular basis of A-T is loss of ATM protein. The majority of A-T missense mutations are found within the FAT-KIN domains (Choi et al., 2016) and ATM is often mutated in multiple cancers, with the FAT-KIN regions exhibiting some of the highest mutational frequency (Forbes et al., 2015; McKusick, 2007). Consistent with loss of activity in A-T and possibly in some cancers, some of the cancer-associated mutations are present in the highly conserved glycine-rich loop, the catalytic loop as well as the activation loop, thus affecting catalytic activity directly. A good example is the A-T mutation G2867R (G2609 in Tel1), which sits within the catalytic loop at a sharp turn just below the activation loop (**Figure 4D**). The introduction of a large charged side-chain would certainly alter the structure of these elements critical for catalysis.

A large number of disease-associated substitutions are found outside the Kinase active site and are distributed within the FAT domain and Kinase-FAT interface. Strikingly, there is a large cluster of mutations within the FLAP-BE and others around PRD-I, especially in Kα9 and Kα10 and regions surrounding them (**Figure 4, Table S1**). Presumably these mutations affect PRD-I, thus substrate phosphorylation. The AT mutation, V2424G (L2179 in Tel1), sits close to the cancer-associated mutations E2423, R2443Q, E2444K and D2448Φ (A2178, Q2206, R2209, D2210 in Tel1, respectively) (**Figure 4C, Table S1**) and are likely to disrupt the FLAP-BE structure as they hold the two helices together through salt-bridges or hydrophobic interactions. Disrupting the FLAP-BE would likely alter the PRD-I within the context of a dimer and would therefore alter peptide substrate access. Another cluster of mutations exists around LBE and FATC. Mutations in these regions would also affect substrate access by affecting PRD-I. Interestingly, similar to hyperactive mutations, several pathogenic mutations are concentrated around the interfaces between the C-lobe and N-FAT; again these mutations could affect key elements including LBE and the activation loop or prevent allosteric activation.

The nucleotide-bound structure presented here reveals that Tel1 exists as an auto-inhibited dimer that is likely activated via an allosteric mechanism found in other PIKKs. With the exception of N-lobe, the nucleotide co-ordinating residues and the catalytic site in the C-lobe are positioned to allow catalysis. However, the PRD-I competes with peptide-substrates and occludes the access to the active site, and is likely the major determinant of the low basal activity of Tel1. The PRD-I is held in place by a number of PIKK-specific features, with mutations in these regions leading to altered activity and disease phenotypes. The movement of the PRD-I is clearly necessary for full kinase activity and based on other related PIKKs may occur through an allosteric mechanism that is common to this family of kinases. Hyper-activating mutations and disease-associated mutations of Tel1 and ATM clearly suggest the FAT-domain is involved in regulating the activity of this kinase. Further structural work on an activator-bound Tel1^ATM^ will be required to resolve how the PRD-I is evicted from the active site to allow peptide-substrate binding.

## ACKNOWLEDGMENTS

Initial screening of samples was carried out at Imperial College London Centre for Structural Biology EM facility. High-resolution data were collected at the eBIC (proposal EM19865), Diamond Light Source and we thank Drs C. Alistair Seibert and Yuriy Chaban for their support in collecting the data. eBIC is funded by the Wellcome Trust, MRC and BBSRC. This work is funded by the Wellcome Trust Investigator Award to XZ (210658/Z/18/Z), the National Institutes of Health (GM118129 to PB) and a NSF Graduate Research Fellowship (2014157291 to SH).

## AUTHOR CONTRIBUTIONS

XZ and LAY designed the studies. SH and LAY prepared the samples. LAY with RA carried out initial cryoEM studies. LAY and RMW performed the cryoEM analysis, built and refined the structural models. XZ and PB supervised the studies. XZ, LAY and RMW wrote the manuscript with input from all the authors.

## DECLARATION OF INTERESTS

The authors declare no competing interests.

## METHODS

### Electron Microscopy grid preparation

A frozen aliquot of *Saccharomyces cerevisiae* Tel1 (800nM, stored in 40mM HEPES 7.8, 10% Glycerol, 200mM NaCl, 2mM DTT, 0.1% Tween-20, 0.01% NP40, 1mM EDTA, 0.5mM EGTA) was diluted 8-fold using sequential addition of 10 μl volumes of buffer (50mM Tris-HCl, 50mM NaCl, pH 7.4 supplemented with AMP-PNP and Magnesium Acetate), and was incubated for 30 minutes on ice. The final concentration of Tel1 was 100nM; 1mM AMP-PNP; 4mM Mg(OAc)_2_. Samples (4 μl) were deposited onto Lacey Carbon 300 mesh gold grids that also have an additional ultrathin carbon support layer (Ted Pella Inc. USA), which were plasma-cleaned, for 30 seconds in air, prior to sample application. Samples were vitrified in liquid ethane at liquid nitrogen temperature using a Vitrobot Mk IV (FEI) set with a blotting force of -6, a waiting time of 60 s and a blotting time of 2 seconds. Plunge freezing was performed at 4 °C and 100% humidity.

### CryoEM Data Acquisition

High-resolution data were collected for Tel1 over three sessions at eBIC (Oxfordshire, UK) on an FEI Titan KRIOS (Thermo Fisher) and are summarised in Table 1. For all three datasets, the microscopes were operated at 300kV with the specimen at cryogenic temperatures (approximately -180 °C) with images recorded at 1-3 μm underfocus on a Falcon III direct electron detector in linear mode at a nominal magnification of 75,000 X, corresponding to a calibrated pixel size of 1.09 Å, and a cumulative total electron dose of ∼90 e^-^/Å^2^. We collected a total of 12,609 micrographs, which were fractionated into frames (dataset 1, 4508, 23 frames; dataset 2; 2445, 34 frames; dataset 3; 5656, 34 frames). A representative micrograph is shown in Figure S1A.

**Table 1.**
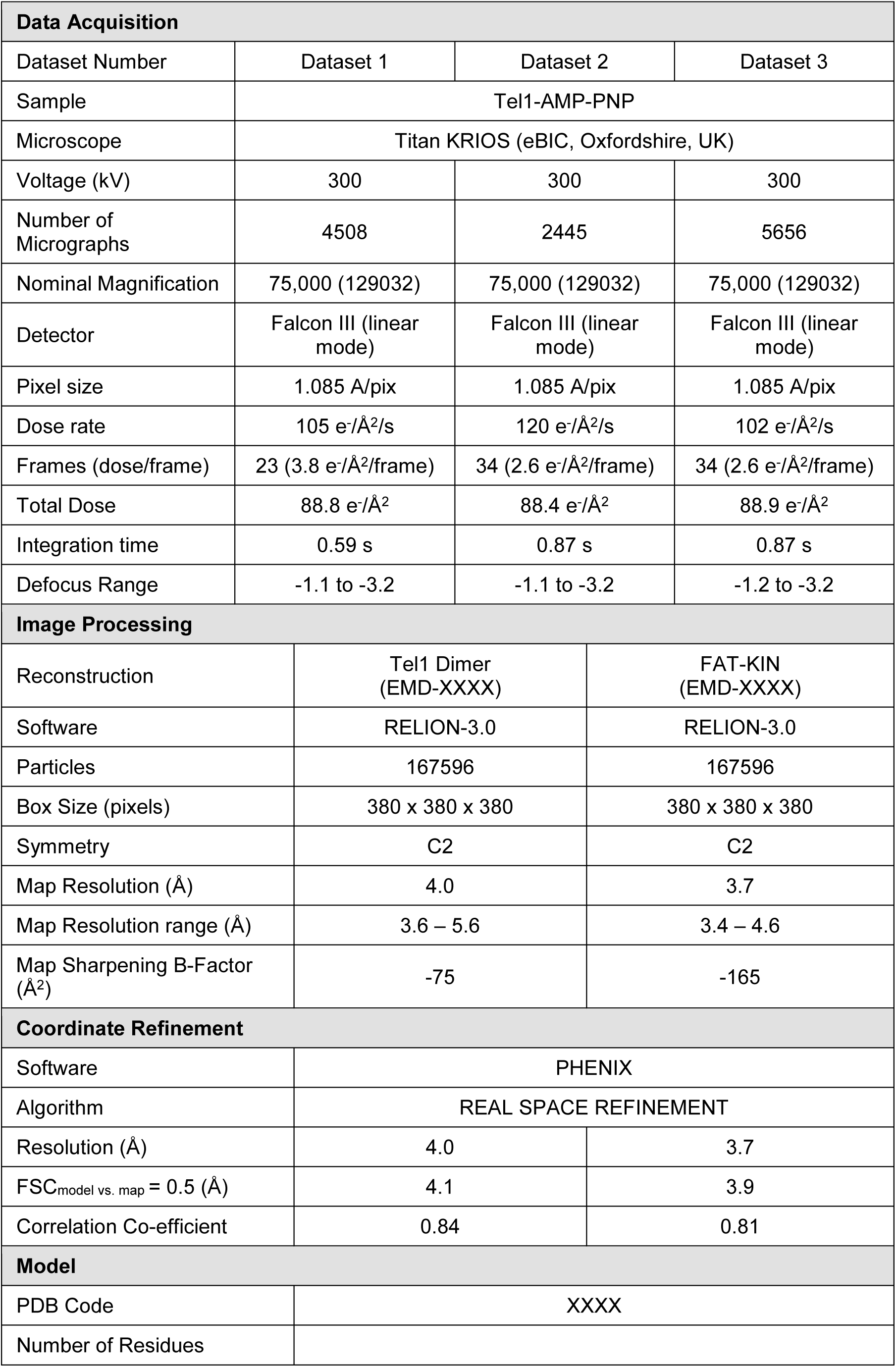
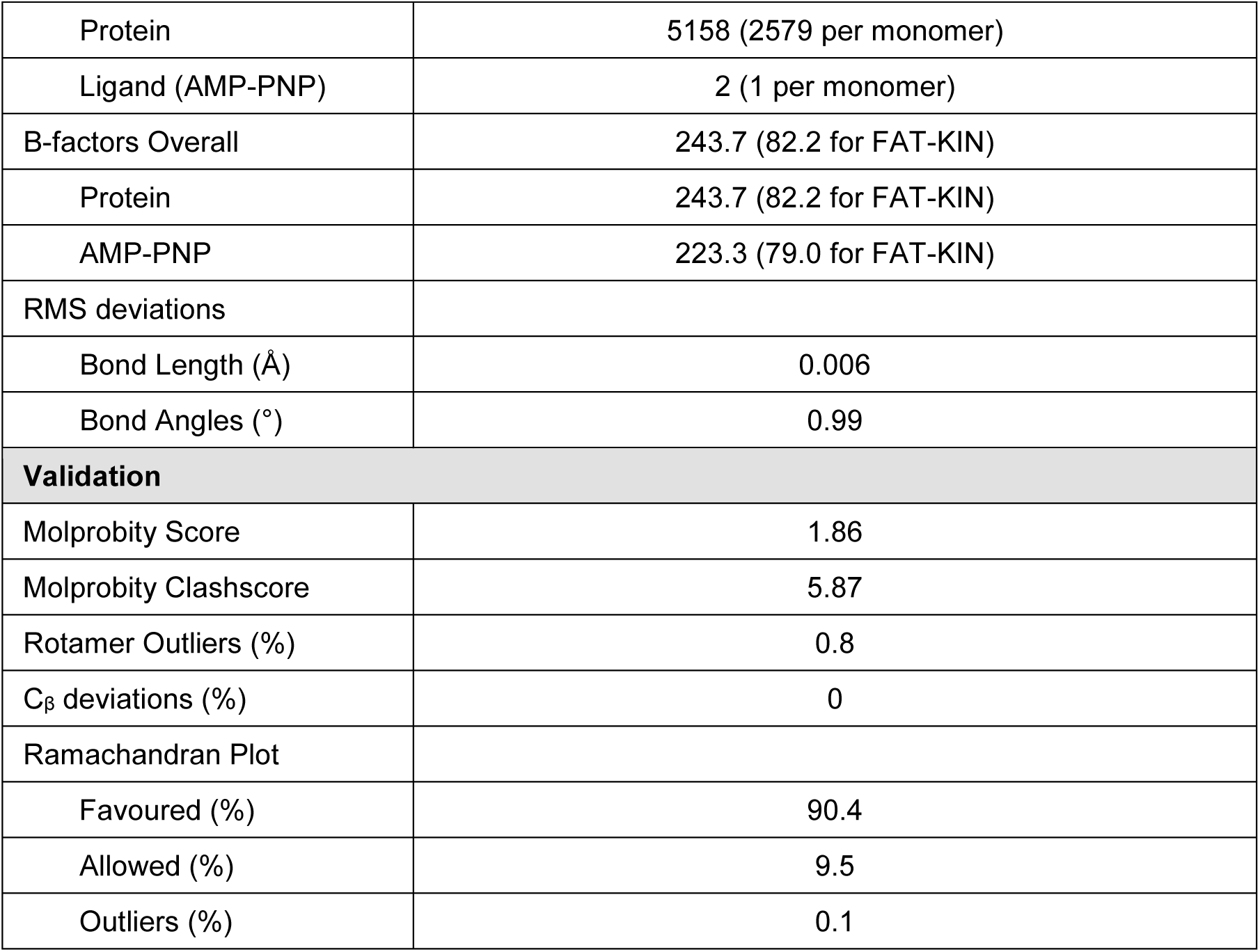
Data collection and Refinement statistics.

| Data Acquisition |  |  |  |
| --- | --- | --- | --- |
| Dataset Number | Dataset 1 | Dataset 2 | Dataset 3 |
| Sample | Tel1-AMP-PNP |  |  |
| Microscope | Titan KRIOS (eBIC, Oxfordshire, UK) |  |  |
| Voltage (kV) | 300 | 300 | 300 |
| Number of Micrographs | 4508 | 2445 | 5656 |
| Nominal Magnification | 75,000 (129032) | 75,000 (129032) | 75,000 (129032) |
| Detector | Falcon III (linear mode) | Falcon III (linear mode) | Falcon III (linear mode) |
| Pixel size | 1.085 Å/pix | 1.085 Å/pix | 1.085 Å/pix |
| Dose rate | 105 e<sup>-</sup>/Å<sup>2</sup>/s | 120 e<sup>-</sup>/Å<sup>2</sup>/s | 102 e<sup>-</sup>/Å<sup>2</sup>/s |
| Frames (dose/frame) | 23 (3.8 e<sup>-</sup>/Å<sup>2</sup>/frame) | 34 (2.6 e<sup>-</sup>/Å<sup>2</sup>/frame) | 34 (2.6 e<sup>-</sup>/Å<sup>2</sup>/frame) |
| Total Dose | 88.8 e<sup>-</sup>/Å<sup>2</sup> | 88.4 e<sup>-</sup>/Å<sup>2</sup> | 88.9 e<sup>-</sup>/Å<sup>2</sup> |
| Integration time | 0.59 s | 0.87 s | 0.87 s |
| Defocus Range | -1.1 to -3.2 | -1.1 to -3.2 | -1.2 to -3.2 |
| Image Processing |  |  |  |
| Reconstruction | Tel1 Dimer (EMD-XXXX) | FAT-KIN (EMD-XXXX) |  |
| Software | RELION-3.0 | RELION-3.0 |  |
| Particles | 167596 | 167596 |  |
| Box Size (pixels) | 380 x 380 x 380 | 380 x 380 x 380 |  |
| Symmetry | C2 | C2 |  |
| Map Resolution (Å) | 4.0 | 3.7 |  |
| Map Resolution range (Å) | 3.6 – 5.6 | 3.4 – 4.6 |  |
| Map Sharpening B-Factor (Å<sup>2</sup>) | -75 | -165 |  |
| Coordinate Refinement |  |  |  |
| Software | PHENIX |  |  |
| Algorithm | REAL SPACE REFINEMENT |  |  |
| Resolution (Å) | 4.0 | 3.7 |  |
| FSC<sub>model vs. map</sub> = 0.5 (Å) | 4.1 | 3.9 |  |
| Correlation Co-efficient | 0.84 | 0.81 |  |
| Model |  |  |  |
| PDB Code | XXXX |  |  |
| Number of Residues |  |  |  |

|  |  |
| --- | --- |
| Protein | 5158 (2579 per monomer) |
| Ligand (AMP-PNP) | 2 (1 per monomer) |
| B-factors Overall | 243.7 (82.2 for FAT-KIN) |
| Protein | 243.7 (82.2 for FAT-KIN) |
| AMP-PNP | 223.3 (79.0 for FAT-KIN) |
| RMS deviations |  |
| Bond Length (Å) | 0.006 |
| Bond Angles (°) | 0.99 |
| **Validation** |  |
| Molprobrity Score | 1.86 |
| Molprobrity Clashscore | 5.87 |
| Rotamer Outliers (%) | 0.8 |
| C<sub>β</sub> deviations (%) | 0 |
| Ramachandran Plot |  |
| Favoured (%) | 90.4 |
| Allowed (%) | 9.5 |
| Outliers (%) | 0.1 |

### CryoEM Image Processing

Movie frames were aligned, corrected for drift, beam induced motion and dose-weighted using MotionCor2 (Zheng et al., 2017) using a 5 × 5 patch implemented in RELION-3.0 (Zivanov et al., 2018). Contrast transfer function (CTF) fitting was performed using Gctf (Zhang, 2016). Particles were picked with Gautomatch using re-projections of a low resolution Tel1 EM structure (Sawicka et al., 2016) for selection. Particles were extracted in RELION-3.0 using a box size of 400 x 400 pixels and binned four times for initial processing. The number of particles picked per dataset are summarised in Table 1. Reference-free 2D classification of particles from a single dataset was performed in RELION-3.0 and revealed two major views (Figure S1A) and subsequent 3D volumes after refinement showed preferential orientation issues. Therefore, 2D classification was omitted as a first step and 3D classification with an initial Tel1 model (Sawicka et al., 2016), filtered to 60 Å, was performed instead for all datasets. Particle star files were sub-divided to speed up the processing pipeline, as the datasets still contained non-particles and ‘junk’ not removed by 2D classification. Initial 3D classification with 4 classes produced a single class (23% particles) that exhibited features expected for this protein. Similar 3D classes from each batch/dataset were selected, re-extracted twice binned, and joined before an initial consensus 3D refinement. After 3D refinement the particle stack was further cleaned using 3D classification in RELION-3.0 using local angular searches, higher T-factors (T=8) and sub-dividing into six classes. Of the 6 classes many displayed over-fitted and noisy features, however a single class (29%, ∼160K particles) showed clear secondary structure and was selected for further refinement. Particles corresponding to the best 3D class were re-extracted (1.085 Å/pixel) in a slightly smaller box size (380 x 380 pixels) and were refined according to the gold-standard refinement procedure implemented in RELION-3.0 applying C2 symmetry and using a soft mask (C2 symmetric) corresponding to the protein, resulting in a 4.4 Å reconstruction after post-processing according to the FSC = 0.143 criterion (Scheres and Chen, 2012). Beam-induced particle polishing (Zivanov et al., 2019) followed by CTF refinement (Zivanov et al., 2018) improved the map resolution to 4.0 Å. Local resolution estimates calculated in RELION-3.0 showed that density regions corresponding to the N-terminal HEAT repeats were of lower resolution (∼5 Å) as compared to the kinase-containing C-terminal half (∼3.6 – 4.4 Å, Figure S2A). Using a soft mask (C2 symmetrized) encompassing some of the HEAT repeat regions, the FAT domain and the C-terminal kinase domain of the dimer, 3D refinement (gold-standard) improved the resolution yielding a 3.7 Å reconstruction after post-processing (according to FSC = 0.143 criterion), corresponding to approximately two thirds of the protein. Angular accuracy and angular distribution plots suggest that rare views were captured by omitting initial 2D classification steps and that the early preferential orientation issues were circumvented. Local resolution estimates show that the best resolution regions are better than 3.5 Å and the map shows defined secondary structure features and clearly resolved bulky side chains (Figure S2). A set of 167596 particles images (∼47% from dataset 1, ∼12% from dataset 2, ∼41% from dataset 3) were used for the final reconstructions.

### Tel1 Model Building and Refinement

The final 3.7 Å and 4.0 Å EM maps were sharpened with a negative B-factor of -165, as determined by RELION-3.0, or less to avoid high-resolution noise and therefore over fitting of a model. A structure of dimeric human ATM (pdb 5np0) was docked into the maps in the first instance. It was clear from this initial fitting that AMP-PNP was bound in the active site. The ATM model, which is restricted to Cα-Cβ only, was manually fitted as rigid body regions into our maps and residues that did not fit the density or that clearly differed between species were trimmed in Coot (Emsley and Cowtan, 2004). The high-resolution maps permitted accurate model building and therefore we built the structure manually starting with the kinase bound to AMP-PNP using the high-resolution nucleotide-bound X-ray structures of mTOR (Yang et al., 2013) and PKA (Das et al., 2015) as guides alongside side-chain density in our maps to model the correct location of the catalytic loop, activation loop and the glycine-rich loop. The kinase domain was subsequently built manually by using residues at the AMP-PNP bound active site as a start point and using bulky side chains as landmarks for correct sequence assignment against UNIPROT accession code P38110. We also made use of a local agreement filtering program (Local Agreement Filtering Algorithm for Transmission EM Reconstructions [LAFTER]) (Ramlaul et al., 2019) that produces a map filtered to maintain consistent features between the two independent half-maps from gold-standard refinement and recovers more signal. This allowed us to confidently place residues in loops and regions where sharpening did not aid model building. We were able to build co-ordinates with the majority of side chains corresponding to residues 1527-2787 and place residues with sequence assignment (but occasional side chains) to 967-1526. The N-HEATs were more challenging to build and so these were built as a poly-Alanine trace. The co-ordinates for the model corresponding to the FAT-Kinase regions of the dimer were first real space refined in PHENIX (Adams et al., 2010; Afonine et al., 2018) against the 3.7 Å map (sharpened with -165 B-factor). The Tel1 Dimer co-ordinates (which include the previously refined FAT-Kinase region) were refined against the 4.0 Å map (sharpened with -75 B-factor). In both cases data used in refinement was limited to spatial frequencies to the RELION estimated resolution to prevent over-fitting. Ramachandran, C_β_, non-crystallographic symmetry (NCS), and secondary structure restraints (generated in PHENIX using caBLAM) were used throughout the refinement to ensure good model geometry and the coordinates were validated using MOLPROBITY (Chen et al., 2015) in PHENIX. Typically 3-cycles of real space refinement were run (3 macro cycles of global and local optimization and B-factor refinement), with PHENIX automatically estimating relative weighting of the restraints and map to prevent over-fitting (Afonine et al., 2018). Refinement and model statistics are given in Table 1. Map vs model FSC curves were also generated in PHENIX as part of the refinement procedure and given in Figure S2.

### Model Interpretation and Analysis

Figures were created using PyMOL (Schrodinger, LLC) and UCSF Chimera (Pettersen et al., 2004). Structural superposition of structures was performed in PyMOL aligning kinases by their C-lobes. Dimer interface buried surface area estimates were calculated using PISA (Krissinel and Henrick, 2007). Multiple sequence alignments were performed using Clustal Omega (Sievers et al., 2011) and displayed in Jalview. Structure-based sequence alignments were performed in PROMALS3D (Pei and Grishin, 2007).

---

**Table S1.**
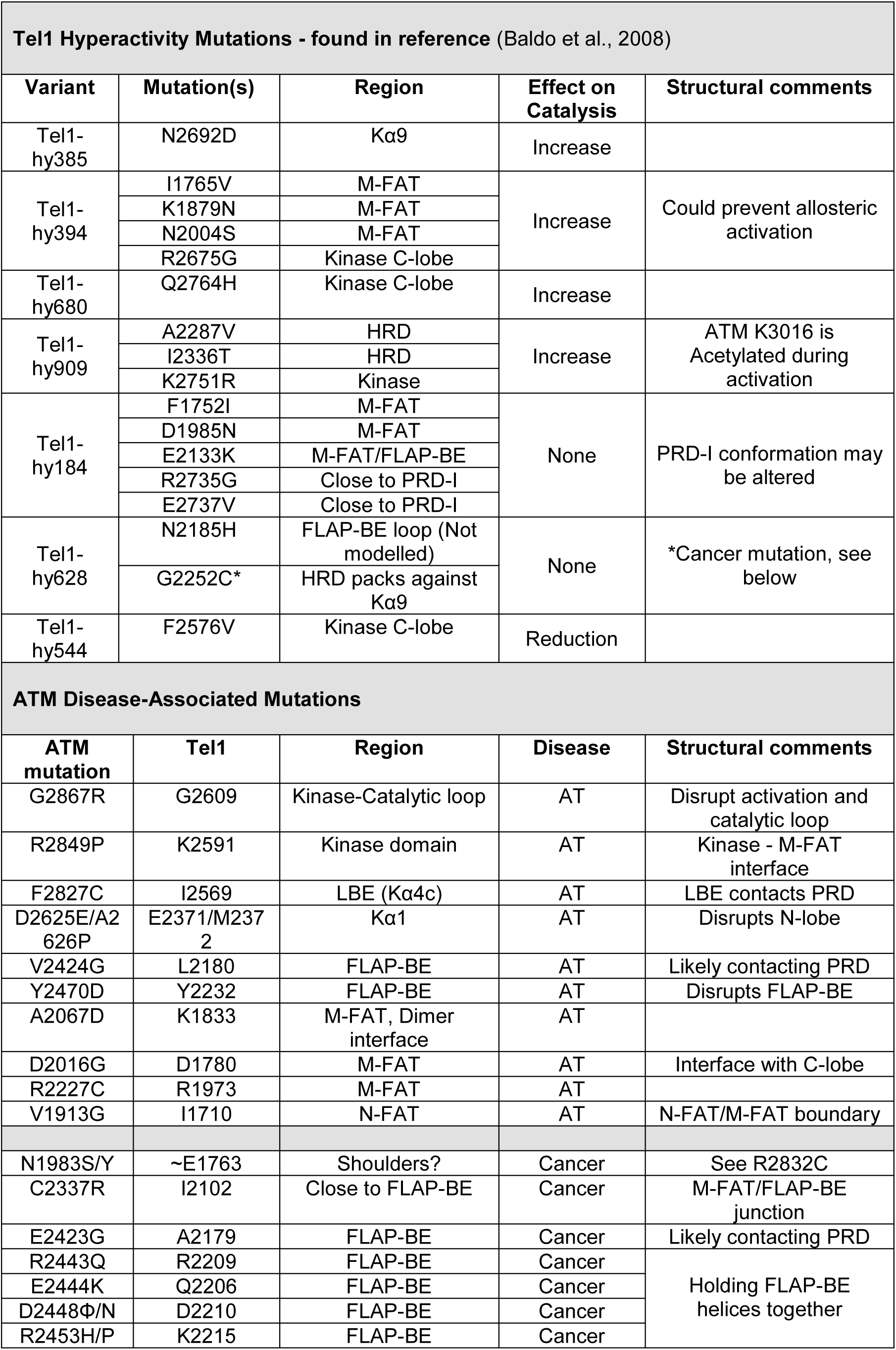
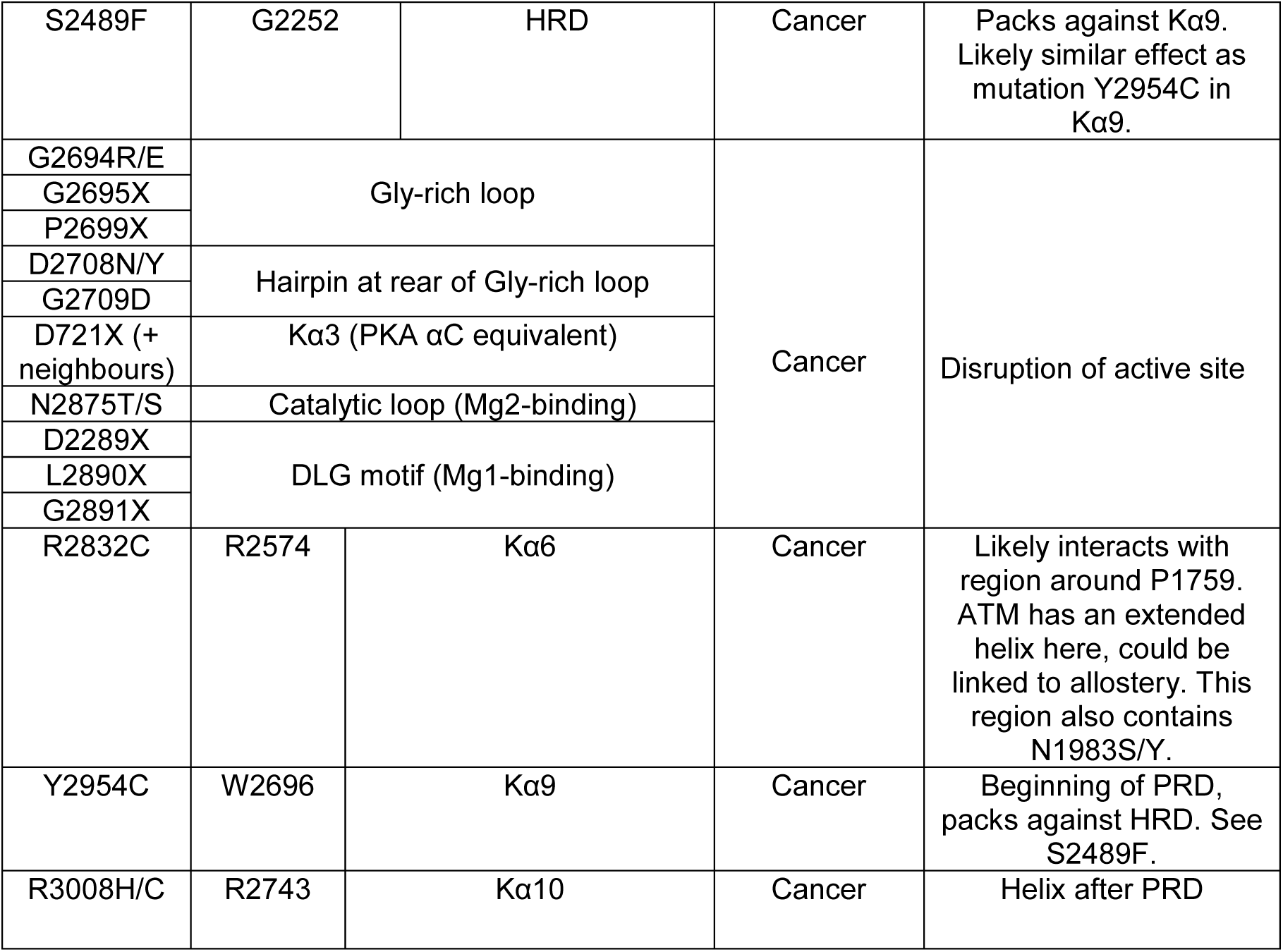

