## Supplemental Figure 1 for "Structure of nucleotide-bound Tel1^ATM^ reveals the molecular basis of inhibition and structural rationale for disease mutations"

**Yates et al. Figure S1**

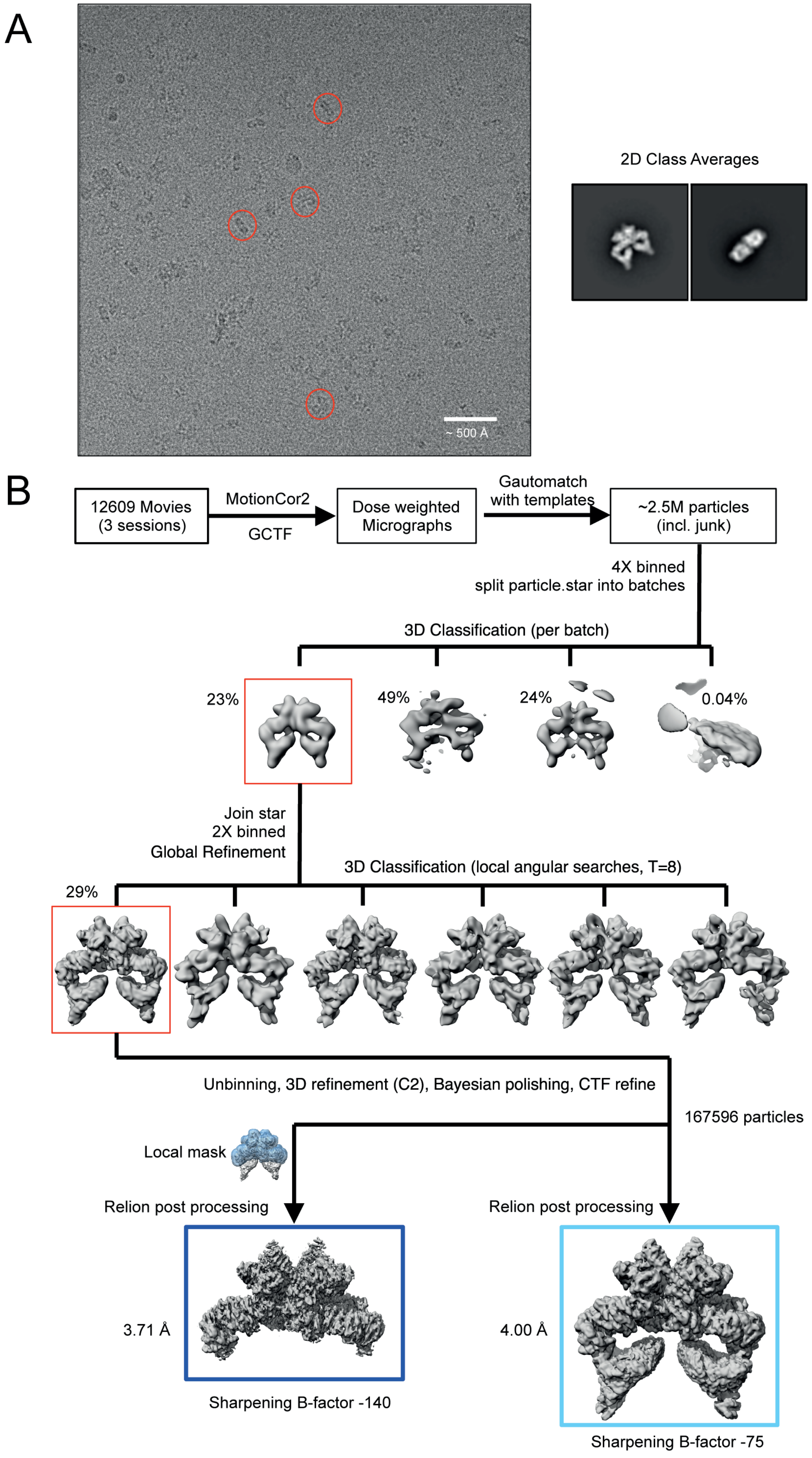


(A) Representative micrograph, with a number of Tel1 particles circled showing distinct views of the protein in the raw image. (B) Summary of the data processing workflow.
