## Supplemental Figure 2 for "Structure of nucleotide-bound Tel1^ATM^ reveals the molecular basis of inhibition and structural rationale for disease mutations"

**Yates et al. Figure S2**

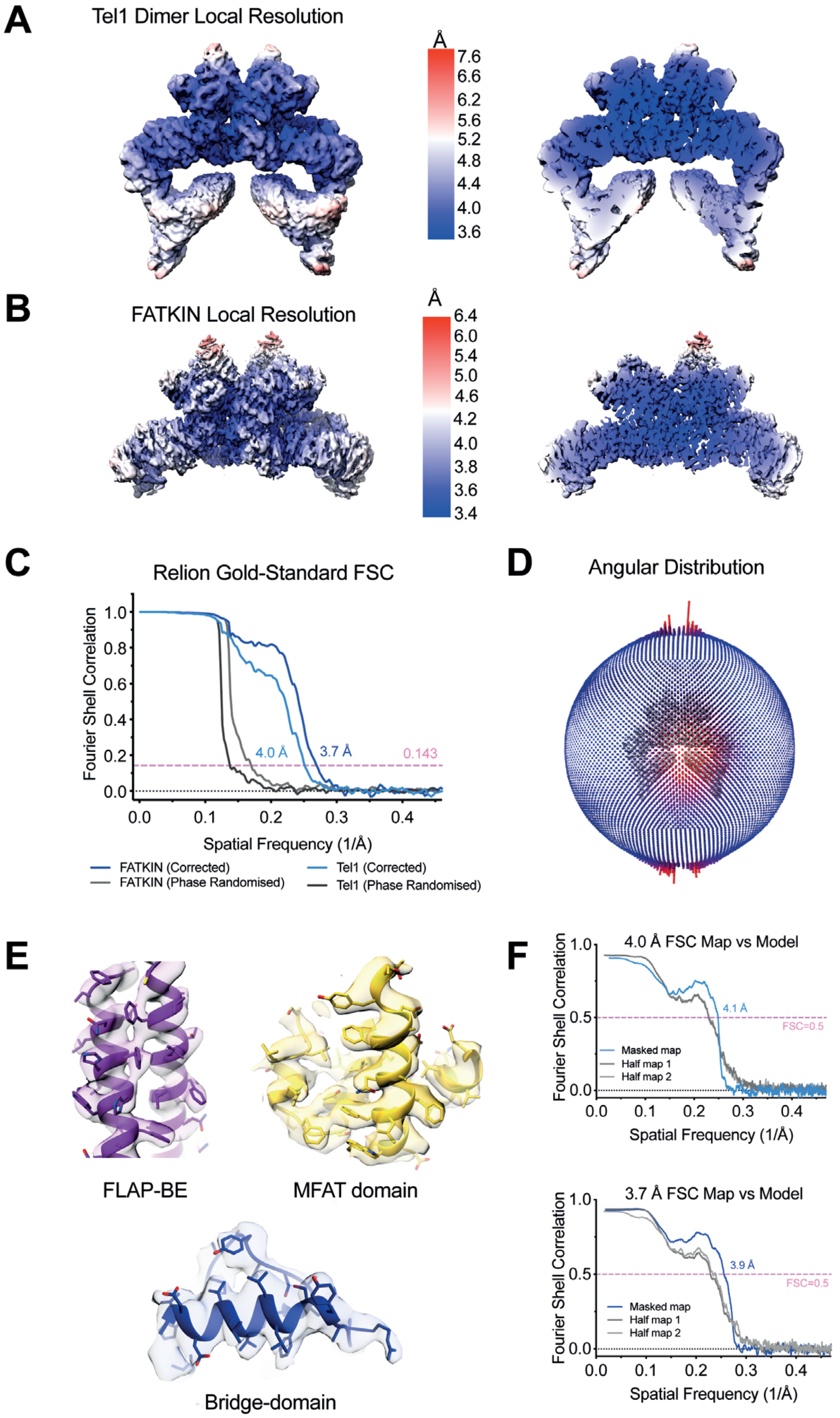


Local resolution estimates of the (A) Tel1 dimer reconstruction and (B) FAT-KIN reconstruction calculated in RELION-3.0. A cut-away view into the core of the reconstruction is also shown. (C) Corrected Fourier Shell Correlation (FSC) curves. (D) Angular distribution of the Tel1 dimer reconstruction. (E) Examples of density regions showing clear secondary structure and side chain details. (F) Map versus model FSC after real space refinement of the Tel1 coordinates into the maps.
