## Supplemental Figure 3 for "Structure of nucleotide-bound Tel1^ATM^ reveals the molecular basis of inhibition and structural rationale for disease mutations"

**Yates et al. Figure S3**

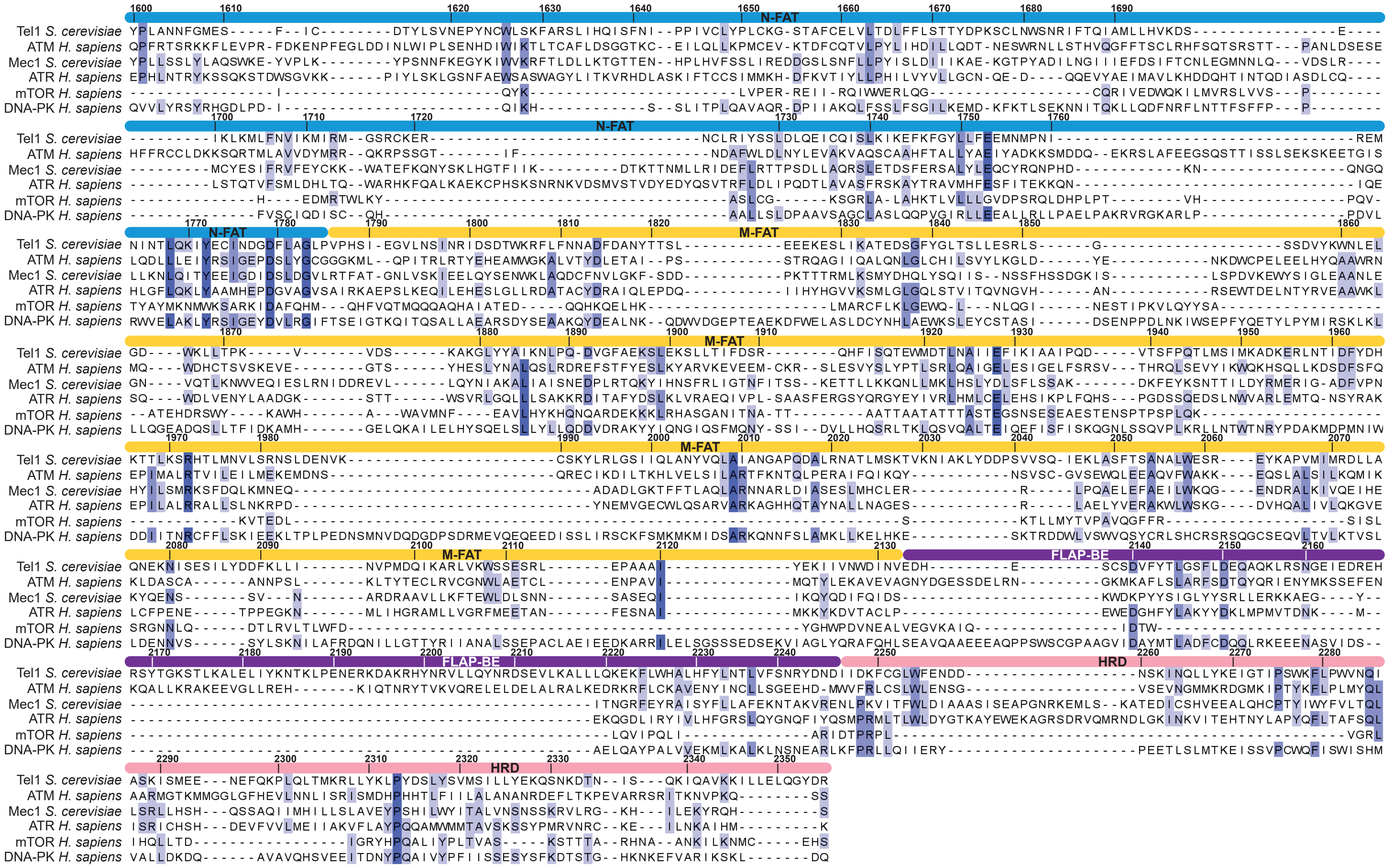


Structure-based multiple sequence alignment of the FAT region of PIKKs. Tel1 residue numbers are shown above the sequence, with domain boundaries coloured as in figure 1.
