## Supplemental Figure 4 for "Structure of nucleotide-bound Tel1^ATM^ reveals the molecular basis of inhibition and structural rationale for disease mutations"

**Yates et al. Figure S4**

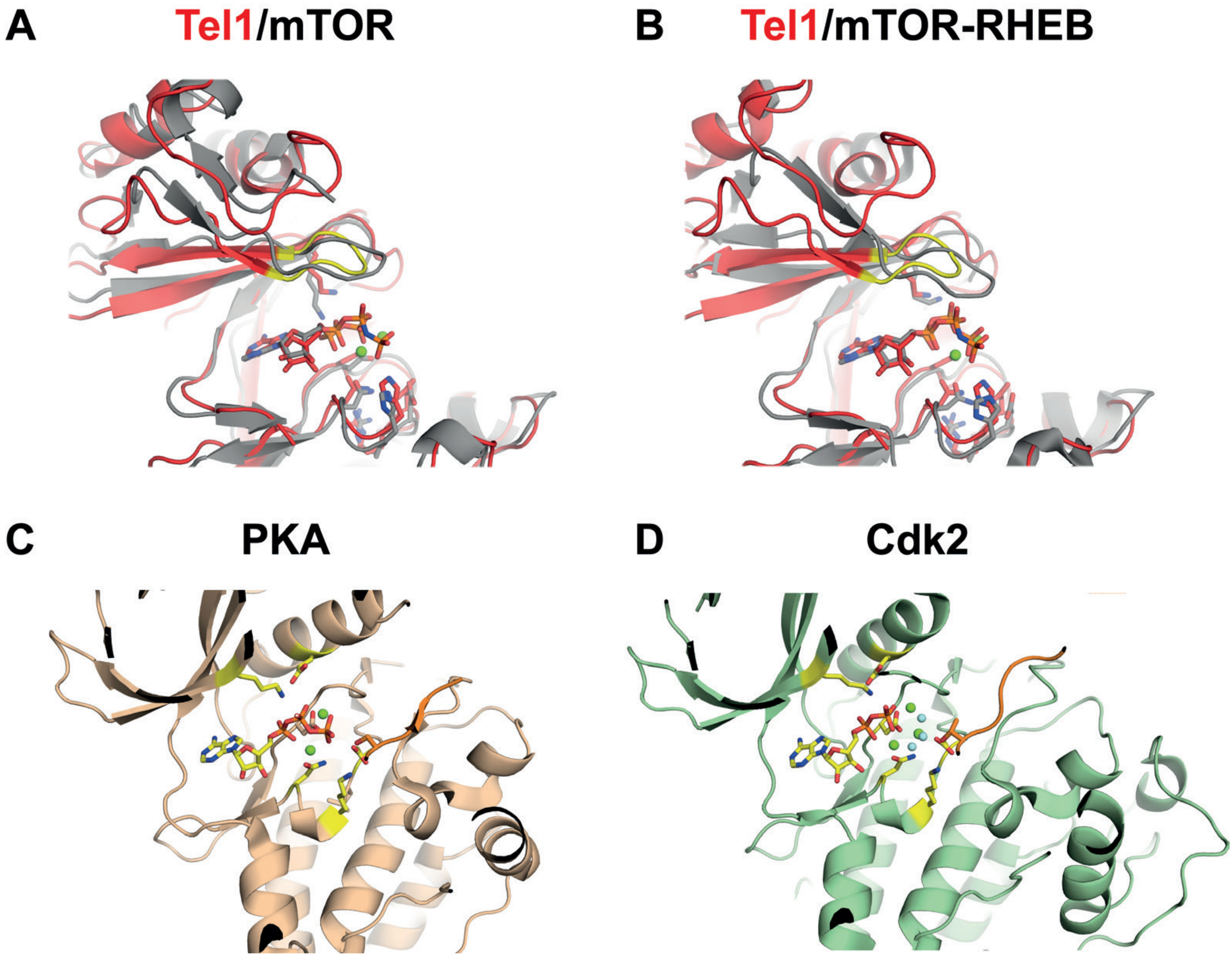


Active site comparisons between (A) Tel1 (red) with mTOR (grey); (B) Tel1 (red) RHEB-activated mTOR (grey). The active sites of two well-described kinases-peptide substrate complexes (C) PKA and (D) Cdk2 are shown for comparison.
