## Supplemental Figure 5 for "Structure of nucleotide-bound Tel1^ATM^ reveals the molecular basis of inhibition and structural rationale for disease mutations"

Yates et al. Figure S5

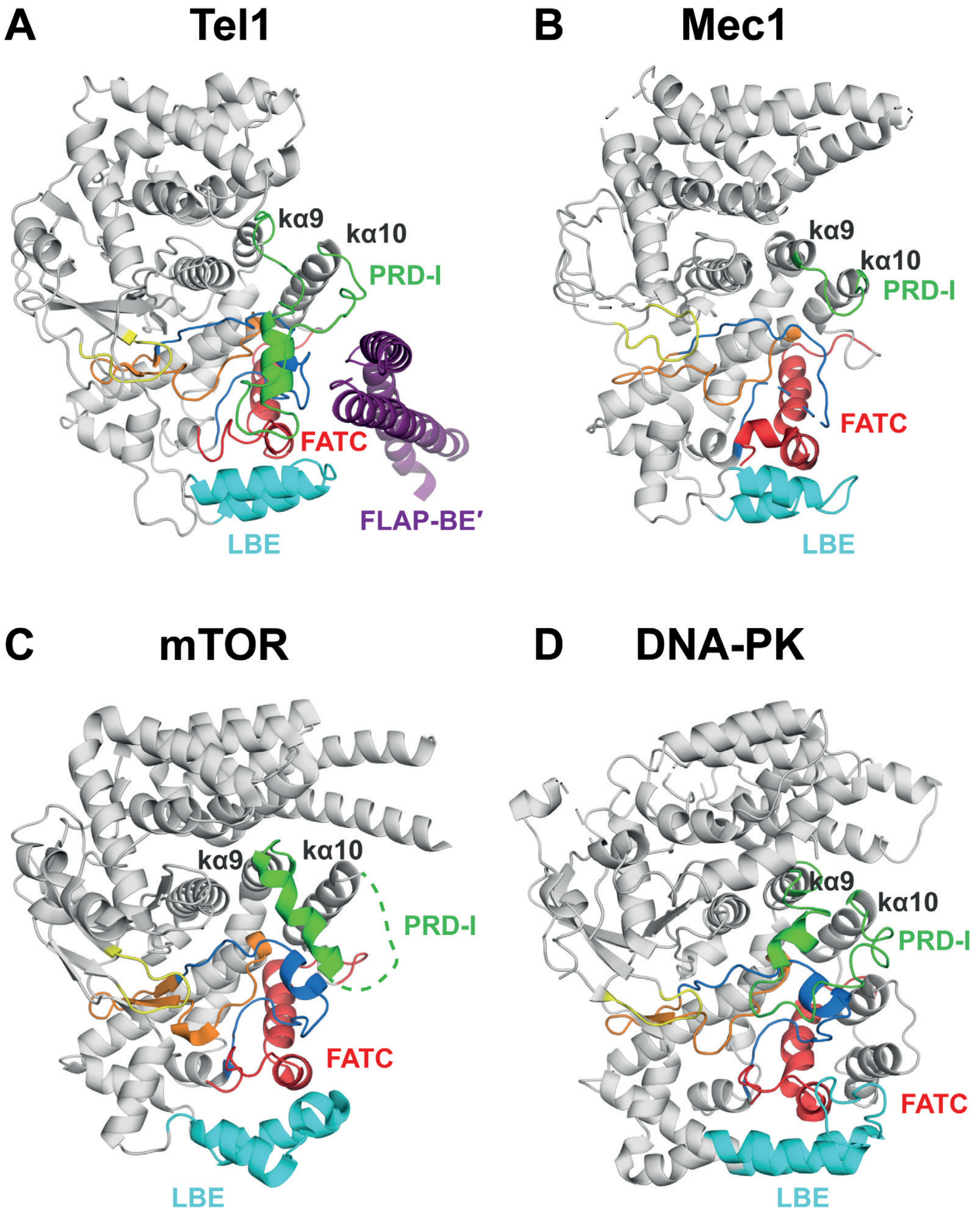


Structural comparisons between the PIKKs (A) Tel1, (B) Mec1, (C) mTOR, and (D) DNA-PK, showing structural divergence in the PRD-I, but conserved features in the catalytic sites, which are coloured as in Figure 3.
