## Supplemental Figure 6 for "Structure of nucleotide-bound Tel1^ATM^ reveals the molecular basis of inhibition and structural rationale for disease mutations"

Yates et al. Figure S6

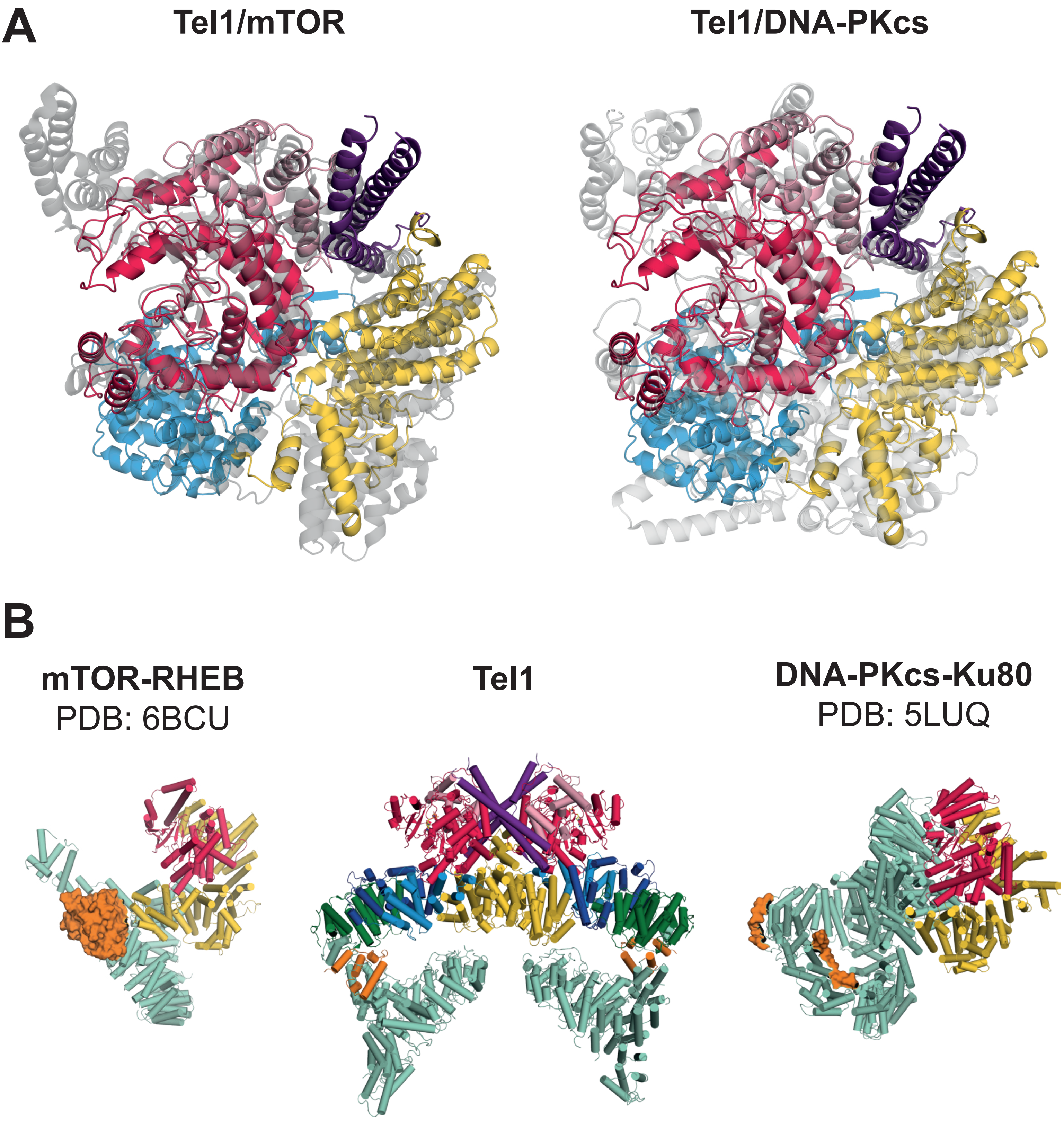


Structural comparison of FAT-KIN regions between (A) Tel1 (coloured as in Figure 1) versus mTOR and Tel1 versus DNA-PK showing a conserved domain arrangement. (B) Activator-binding in mTORC1 (RHEB binding) and DNA-PK (Ku70/80), where activators are rendered orange. Tel1 activator MRX binding site is unknown, but Xrs2 is shown to interact at the approximate region in Tel1, highlighted in orange. mTOR and DNA-PK are coloured by kinase (red), FAT (yellow) and HEAT repeats (green). All structures are aligned on the kinase domain.
